# An essential ER-resident N-acetyltransferase in *Plasmodium falciparum*

**DOI:** 10.1101/2021.11.01.466802

**Authors:** Alexander J Polino, Katherine Floyd, Yolotzin Avila-Cruz, Yujuan Yang, Daniel E. Goldberg

## Abstract

N-terminal acetylation is a common eukaryotic protein modification that involves the addition of an acetyl group to the N-terminus of a polypeptide. This modification is largely performed by cytosolic N-terminal acetyltransferases (NATs). Most associate with the ribosome, acetylating nascent polypeptides co-translationally. In the malaria parasite *Plasmodium falciparum*, exported effectors are translated into the ER, processed by the aspartic protease plasmepsin V and then N-acetylated, despite having no clear access to cytosolic NATs. Here, we used post-transcriptional knockdown to investigate the most obvious candidate, Pf3D7_1437000. We found that it co-localizes with the ER-resident plasmepsin V and is required for parasite growth. However, depletion of Pf3D7_1437000 had no effect on protein export or acetylation of the exported proteins HRP2 and HRP3. Pf3D7_1437000-depleted parasites arrested later in their development cycle than export-blocked parasites, suggesting the protein’s essential role is distinct from protein export.

## Introduction

N-terminal acetylation is among the most common modifications to eukaryotic proteins (Deng and Marmorstein, 2020). Typically during or soon-after translation, an acetyl group is transferred from acetyl-CoA to the N-terminus of a polypeptide. This alteration blunts the N-terminal charge, changing its chemical properties, and altering the way the polypeptide interacts with various biological systems. In some cases, N-terminal acetylation changes a protein’s half-life (Hwang et al., 2010; Park et al., 2015). In others, proper interaction with binding partners relies on acetylation (Scott et al., 2011). Acetylation is performed by N-acetyltransferases (NATs), nearly all of which are cytosolic enzymes that typically associate with the ribosome. In eukaryotes, eight currently known NAT complexes combine to acetylate most cytosolic N-termini, as well as some proteins in the chloroplast lumen.

The malaria parasite *Plasmodium falciparum* marks effectors for export into the host cell with a pentameric amino acid sequence called the *<u>Pl</u>asmodium* <u>ex</u>port <u>el</u>ement (PEXEL) (Hiller et al., 2004; Marti et al., 2004). PEXEL-containing proteins are translated into the parasite endoplasmic reticulum (ER), where PEXEL is cleaved by the aspartic protease plasmepsin V (PM V) (Boddey et al., 2010; Russo et al., 2010) after a conserved leucine. Previous work with exported reporters revealed that following PEXEL cleavage in the ER, the new N-terminus is somehow acetylated (Boddey et al., 2009; Chang et al., 2008; Osborne et al., 2010). This acetylation occurred even if exported proteins were sequestered in the ER by brefeldin A treatment (blocking anterograde traffic from the ER) or addition of an ER retention signal (Chang et al., 2008; Osborne et al., 2010). This suggests that a yet unidentified NAT exists in the *P. falciparum* ER. Subsequently Tarr, et al. (2013) mutagenized the PEXEL motif of the exported protein REX3 and found that point mutants that were not acetylated also were not exported (Tarr et al., 2013), demonstrating a coincidence between these two processes. The identity of the PEXEL NAT, and its role in export, if any, remains to be determined.

Here, we searched the genome for putative NATs in *P. falciparum* that might access the secretory system. We identified one, Pf3D7_1437000, as a likely candidate for follow-up investigation. Depletion of Pf3D7_1437000 arrested parasite growth in culture, but had no detectable effect on protein export or on exported protein N-acetylation. The phenotype manifested by Pf3D7_1437000 depletion is distinct from that seen after disruption of an essential component of the export pathway, suggesting that the essential role of Pf3D7_1437000 is not in facilitating protein export.

## Results

To search for an ER-resident NAT, we used PlasmoDB to identify *P. falciparum* genes whose sequence was annotated to contain a motif assigning them to the GNAT enzyme superfamily (Aurrecoechea et al., 2009). This search yielded eight genes. Two– Pf3D7_ 0823300 and Pf3D7_0629000 – have been previously studied in *P. falciparum* and are proposed to act as a GCN5 histone acetyltransferase and a glucosamine-phosphate N-acetyltransferase respectively (Cova et al., 2018; Fan et al., 2004). Five of the remaining six have orthologs in metazoans. Pf3D7_1020700 appears similar to the human NAT10, an RNA cytidine acetyltransferase (Ito et al., 2014). Pf3D7_1003300, Pf3D7_0109500, Pf3D7_0805400, and Pf3D7_1323300 are the closest orthologs of the cytosolic human N-acetyltransferases Naa10, Naa20, Naa30, and Naa40 respectively (Fig. 1A) (Chen et al., 2006). The remaining candidate NAT, Pf3D7_1437000, appears to have orthologs among the Apicomplexa, but no obvious representative outside the phylum. None of the candidate NATs has an apparent signal or retention sequence to drive ER localization. However, Pf3D7_1437000 has two predicted transmembrane domains (Fig. 1B), making it the likeliest candidate to access the secretory system. With that in mind, we focused our efforts on characterizing Pf3D7_1437000.

**Figure 1.**
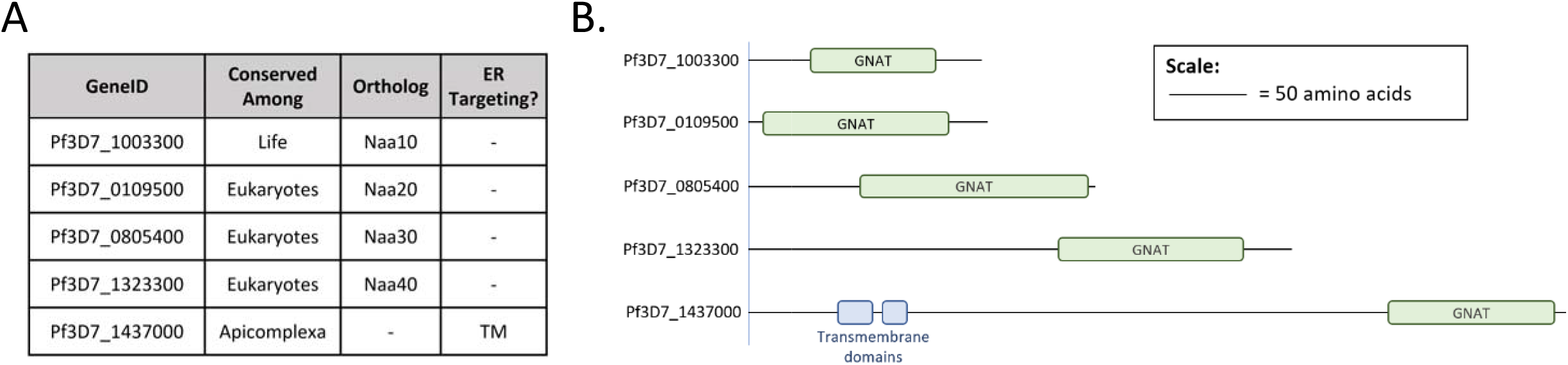
Putative protein N-acetyltransferases encoded by *P. falciparum:* (A) Table of putative N-acetyltransferases (NATs) in the *P. falciparum* genome. Genes were included based on assignment to the GNAT enzyme superfamily in PlasmoDB, and subsequently excluded if they had a different predicted function (see text). Orthology was based on grouping in OrthoMCL, and supported by reciprocal BLAST searches. ER targeting sequences were sought with SignalP 4.1, and transmembrane domains (TM) annotated based on TMHMM predictions. (B) Diagram of the identifiable domains in the NAT candidates in (A). Diagram is to scale as shown.

We targeted Pf3D7_1437000 using CRISPR/Cas9 editing and the previously described pSN054 vector to replace the gene with a recodonized version that is C-terminally HA-tagged and flanked with loxP sites for gene excision, as well as TetR-binding aptamers for post-transcriptional depletion (Fig. 2A) (Polino et al., 2020). Proper genome editing was confirmed by Southern blot (Supp. Fig. 1). Western blotting for HA revealed a single band consistent with tagged full-length Pf3D7_1437000 (expected size 64 kDa) (Fig. 2B). Washing out aTc depleted Pf3D7_1437000 levels to approximately 40% of the +aTc levels within twelve hours (Fig. 2C). Washing out aTc in ring-stage parasites resulted in parasite death within a single intraerythrocytic development cycle, showing Pf3D7_1437000 to be essential for growth in parasite culture (Fig. 2D).

**Figure 2.**
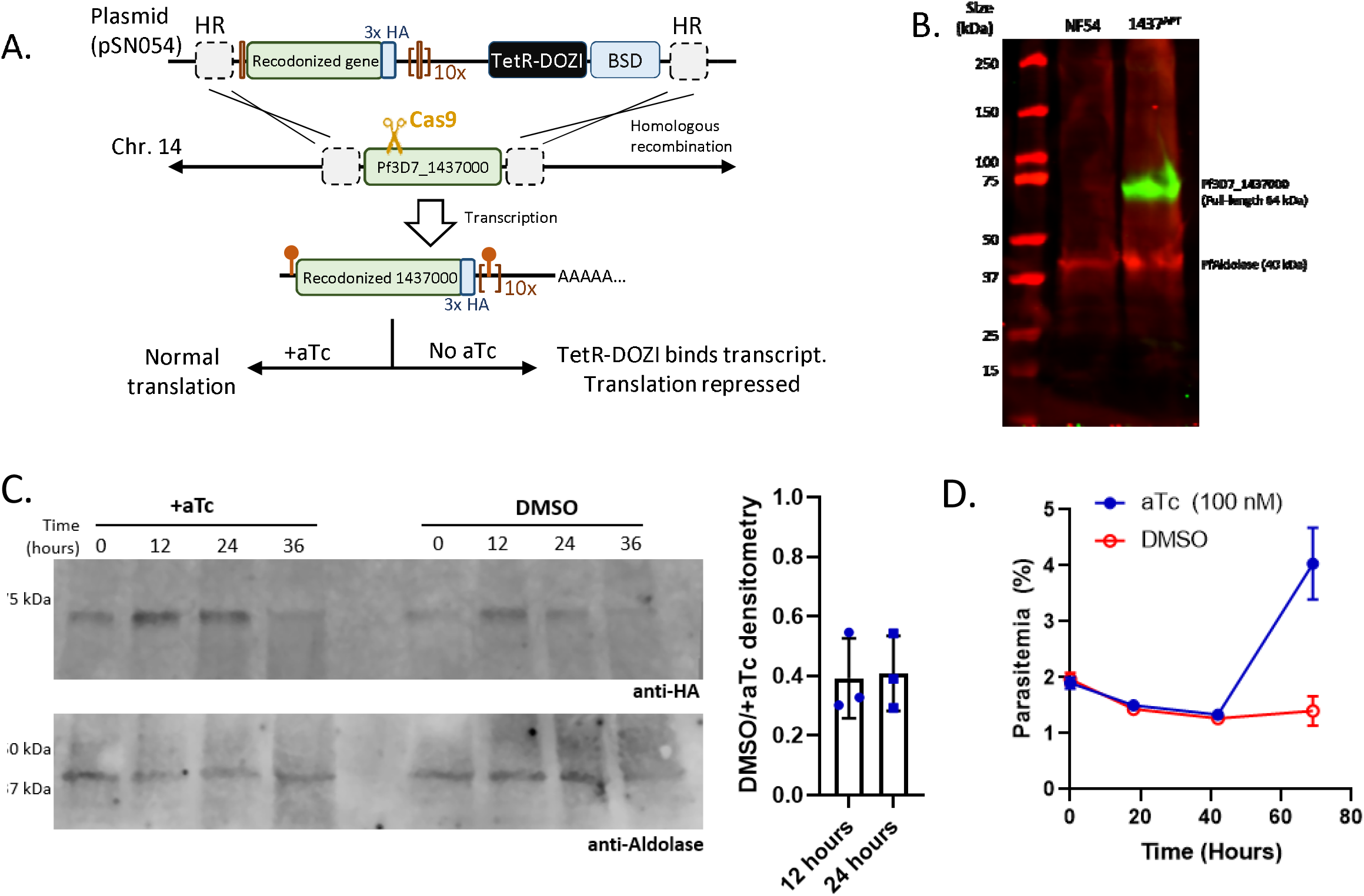
TetR-Aptamer system for inducible Pf3D7_1437000 depletion: (A) Diagram of genome editing and inducible depletion of Pf3D7_1437000. The endogenous gene was disrupted by Cas9 and replaced by a recodonized version flanked with aptamers. When transcribed, the aptamers fold to bind TetR. In the absence of anhydrotetracycline (aTc), TetR binds the aptamers; its fusion protein DOZI sequesters the bound mRNA, repressing translation. (B) Western blot probing lysate from the parent (NF54) or tagged line (1437^APT^) with anti-HA (green) and anti-*P. falciparum* fructose-bisphosphate aldolase (PfAldolase, red). (C) Western blot demonstrating Pf3D7_1437000-HA levels over part of the intraerythrocytic development cycle, and the effect of washing out aTc (“DMSO” as aTc was replaced with an equal volume of DMSO) on Pf3D7_1437000-HA levels. At right, knockdown was quantified for three separate experiments. Bar height represents mean; error bars standard deviation. (D) aTc was washed from ring-stage parasites and their growth monitored daily by flow cytometry. The experiment was performed three times, with each culture in technical triplicate. A representative experiment is shown: points represent the mean of technical triplicates, error bars the standard deviation of those measurements. Uncut gel for panel C is shown in Supp. Fig. 2.

### Pf3D7_1437000 localizes to the ER

We assessed the localization of Pf3D7_1437000-HA in fixed parasites by immunofluorescence. Staining with anti-HA antibodies suggests that Pf3D7_1437000-HA is found in a perinuclear ring, consistent with ER localization (Fig. 3A) (Klemba et al., 2004). This is supported by co-staining with antibodies against organellar markers: anti-plasmepsin V for the ER (Banerjee et al., 2002; Klemba and Goldberg, 2005), anti-ACP for the apicoplast (Waller et al., 1998), and anti-aldolase for the cytosol. The staining pattern of anti-PM V largely resembles that of anti-HA, while anti-ACP and anti-aldolase clearly highlight patterns distinct from anti-HA. To test whether this localization is an artifact of cell fixation, we constructed a parasite line with Pf3D7_1437000 fused to the fluorescent protein mNeonGreen and 3xHA, and PM V fused to mRuby3 and 3xFLAG. Western blot showed most tagged protein to be at the expected size for full-length Pf3D7_1437000-mNeonGreen-3xHA and PM V-mRuby3-3xFLAG (Supp. Fig. 3). Live microscopy on trophozoites showed colocalization between Pf3D7_1437000-mNeonGreen and PM V-mRuby3 (Fig. 3B, C), again suggesting that Pf3D7_1437000 localizes to the parasite ER. Interestingly, we note that both by immunofluorescence and live fluorescence Pf3D7_1437000 and PM V appear to occupy the same space, but their intensities are not perfectly correlated throughout that space: PM V is present in a ring around the nucleus with a single protruding bleb. Pf3D7_1437000 is present over the same space, but disproportionately concentrated in the bleb. The significance of this distinction is not obvious to us.

**Figure 3.**
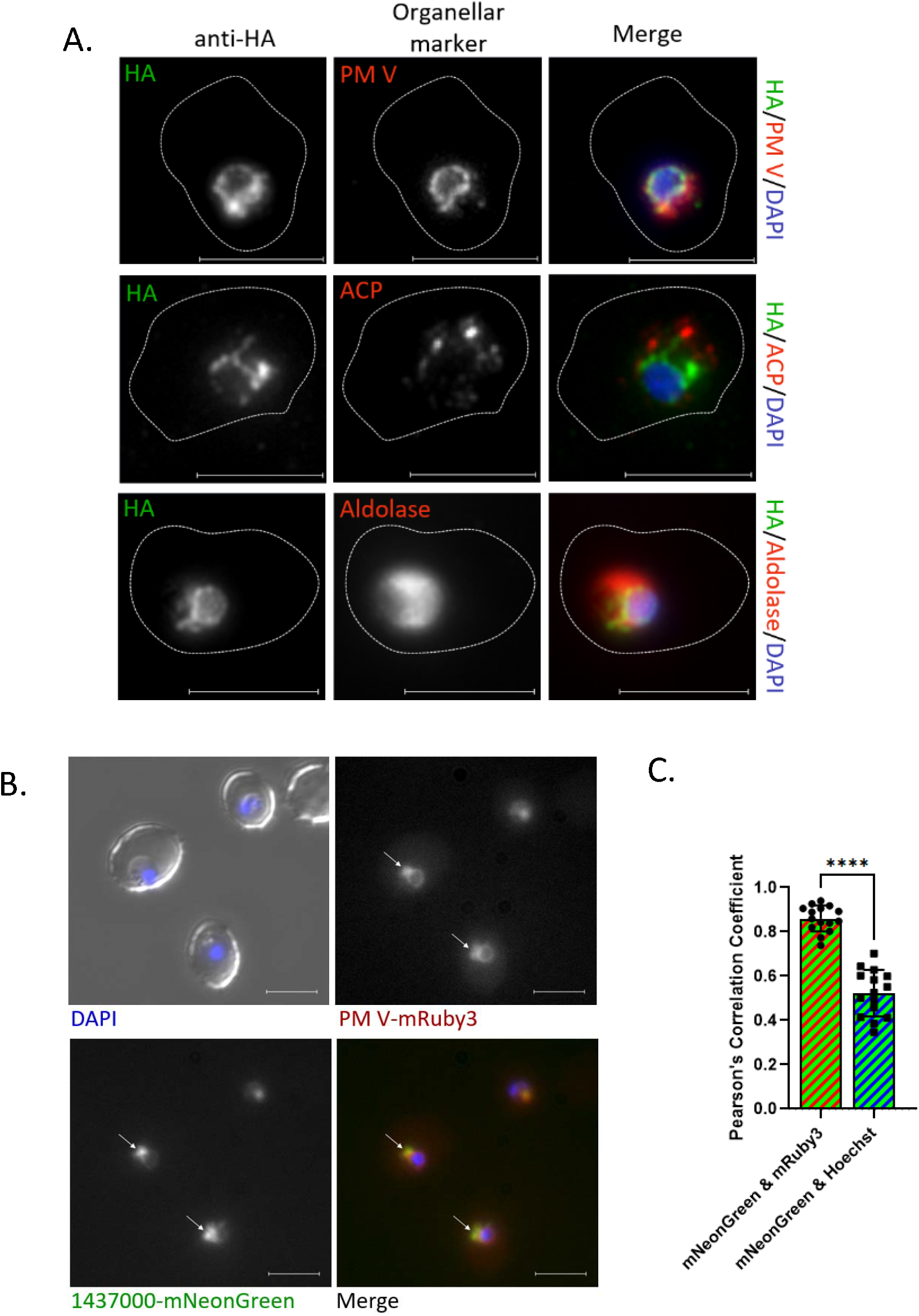
Pf3D7_1437000 resides in the ER: (A) Micrographs of immunofluorescence assay to localize Pf3D7_1437000-HA (green in merge) alongside PM V, ACP, or Aldolase (red in merge). Experiment was performed three times; representative images are shown. (B) Live epifluorescence microscopy with Pf3D7_1437000-mNeonGreen (green) and PM V-mRuby3 (red). Scale bar represents 5 μm. White arrows are to give the reader spatial references between pictures. A representative image is displayed here. Additional zoomed out images are in Supp. Fig. 3. (C) Quantification of signal overlap from (B) by Pearson’s Correlation Coefficient. N = 15 cells. Groups were compared by an unpaired t-test, with p < 0.0001, designated by “****”.

### Pf3D7_1437000 is not required for protein export

Since our original interest was in the acetylation of exported proteins, we next sought to assess whether Pf3D7_1437000 had a role in exporting proteins into the host red blood cell. We used a previously described line where the PTEX component Hsp101 is fused to a DHFR destabilization domain (abbreviated Hsp101-DD), and its function requires the stabilizing ligand trimethoprim (TMP). When TMP is removed, Hsp101 is destabilized and exported proteins accumulate in the parasitophorous vacuole (Beck et al., 2014). Alongside this line, we depleted Pf3D7_1437000 by washing out aTc from young ring-stage parasites (0-4 hours old), then fixed trophozoites and stained for exported proteins by immunofluorescence assay (Fig. 4A). We found that while Hsp101 destabilization blocked protein export as previously described, depletion of Pf3D7_1437000 had no discernible effect on export of the PEXEL proteins HRP II or FIKK4.2 (Fig. 4A, B).

**Figure 4.**
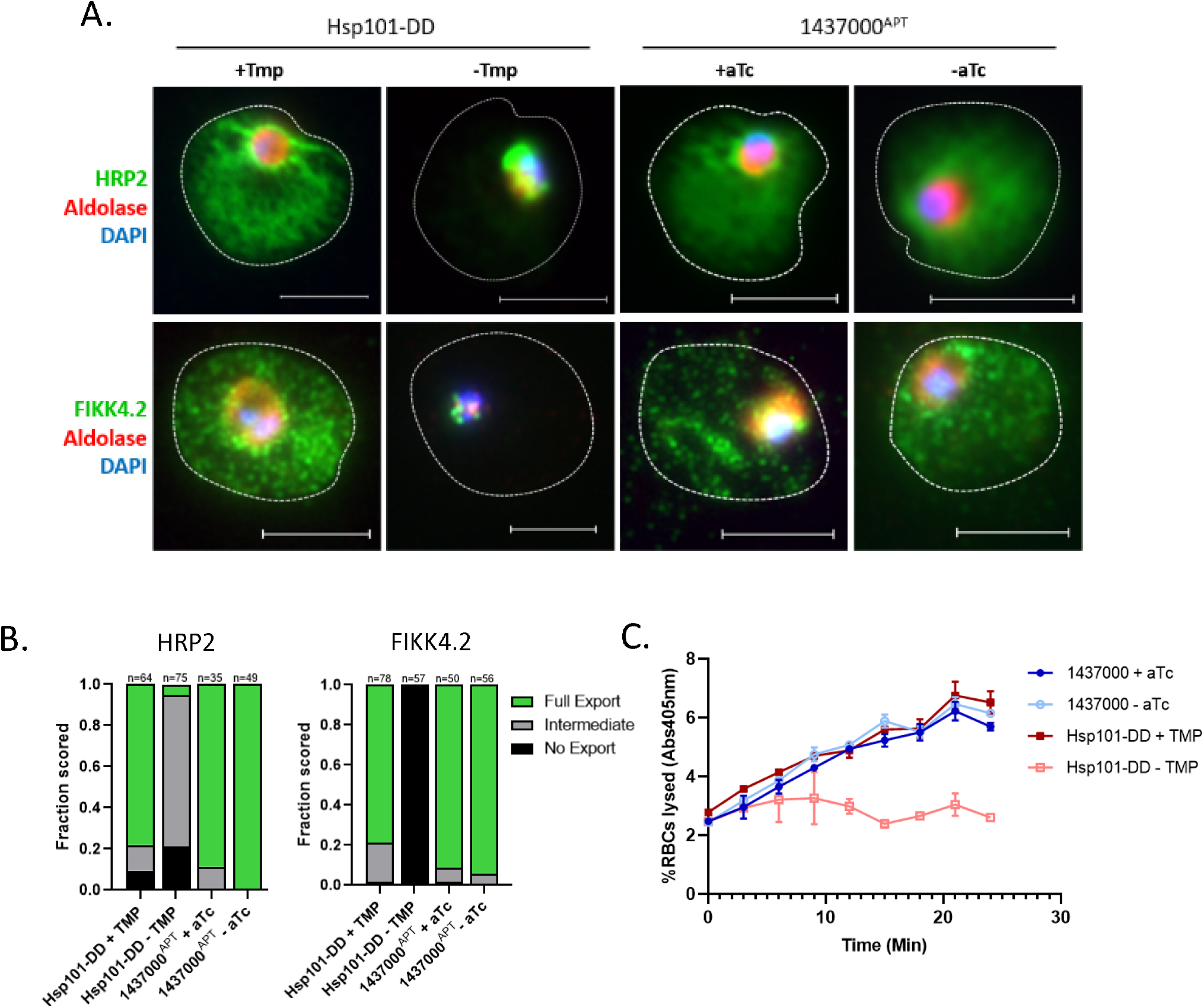
Pf3D7-1437000 depletion does not affect protein export: (A) Trophozoites were fixed and stained to compare protein export in Hsp101-DD +/-TMP and 1437^APT^ +/-aTc. Exported proteins are marked by anti-HRP2 or anti-FIKK4.2, each shown in green; the parasite cytosol is marked by anti-aldolase shown in red; parasite nucleus marked by DAPI in blue. Experiment was performed three times; representative images shown. Scale bars represent 5 μm. (B) Quantification of ten fields by a blinded scorer for the experiment shown in (A). Number of cells counted is shown above each bar. (C) Sorbitol sensitivity was monitored by measuring hemoglobin release into the supernatant (via absorbance at 405 nm) over time. Experiment was performed three times. A representative experiment is shown, with each sample in technical triplicate. Data points represent the measured mean; error bards the standard deviation.

To get an immunofluorescence-independent view of export competence we assessed establishment of the exported protein-dependent nutrient import channel on the infected erythrocyte surface (Beck et al., 2014; Ginsburg et al., 1985). To do so, we depleted Pf3D7_1437000 or disrupted Hsp101, pelleted trophozoites, and resuspended them in 5% sorbitol. Export-competent parasites have increased solute uptake into the host RBC, leaving them susceptible to osmotic lysis in 5% sorbitol. Parasites that cannot export proteins are not able to increase solute uptake, and are protected from sorbitol lysis (Beck et al., 2014; Ginsburg et al., 1985; Kirk et al., 1994). We monitored lysis by measuring the release of hemoglobin into the supernatant over 25 minutes. Disrupting Hsp101 protected parasites from lysis as previously described (Beck et al., 2014) but depleting Pf3D7_1437000 had no effect on sorbitol sensitivity (Fig. 4C), again suggesting Pf3D7_1437000 is not involved in protein export.

To determine whether Pf3D7_1437000 is responsible for acetylating PEXEL proteins, we isolated two abundant exported parasite proteins, histidine-rich proteins II and III (HRP2 and HRP3) (Fig. 5A) and measured the mass of each intact protein by liquid chromatography/ mass spectrometry. In the presence of aTc, we detected substantial peaks consistent with acetylated HRP2 and acetylated HRP3, as expected (Fig. 5B). When we depleted Pf3D7_1437000, we were surprised to find no change in the mass of the peaks (Fig. 5B). In fact, the chromatogram in both cases revealed no peak at the mass expected for unacetylated HRP2 or HRP3.

**Figure 5.**
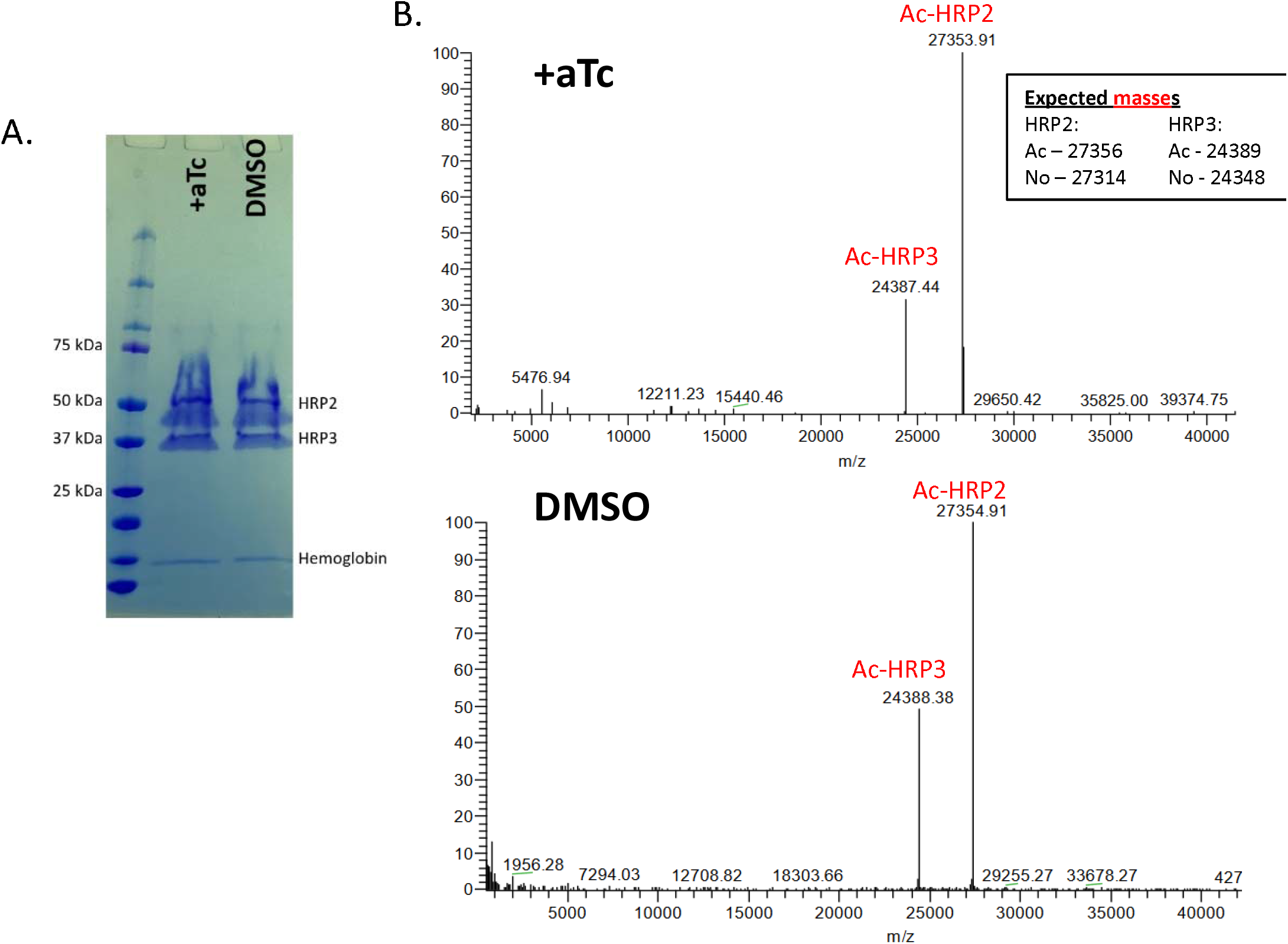
Pf3D7_1437000 depletion does not affect HRP2/HRP3 acetylation: (A) Coomassie-stained SDS-PAGE gel showing HRP2 and HRP3 isolated from the supernatants of saponin-released parasites on nickel resin. Both migrate through the gel less than one would anticipate from their linear size due to their extreme positive charge. (B) Deconvoluted mass spectra from analysis of intact HRP2 and HRP3. Inset shows anticipated sizes for acetylated and un-acetylated HRP2 and HRP3 after PEXEL cleavage.This experiment was performed twice; a representative experiment is shown.

Concerned that this could be due to the slow nature of the aptamer-driven knockdown, we remade the aptamer construct in a parasite line that constitutively expresses a dimerizable Cre recombinase (DiCre) that is activated by the ligand rapamycin (Beck, n.d.; Jullien et al., 2003). Since our aptamer construct already resulted in a gene flanked by loxP sites (Fig. 2A) this allowed us to inducibly excise the Pf3D7_1437000 gene from the genome. Addition of 50 nM rapamycin to the culture media of synchronized ring-stage parasites resulted in complete excision of Pf3D7_1437000 within 24 hours (Supp. Fig. 4A) and parasite death within a single replicative cycle (Supp. Fig. 4B). As before, we purified HRP2 and HRP3 from parasite culture and again found the mass of each unchanged by the excision of Pf3D7_1437000 from the genome (Supp. Fig. 4C). This suggests that Pf3D7_1437000 is not the NAT that acetylates PEXEL proteins.

### Pf3D7_1437000 is involved in parasite growth and entry into schizogony

We next turned our attention to the role of Pf3D7_1437000 in intraerythrocytic development. Normally, parasites are classified as ring forms for the first 20-24 hours after invasion, followed by maturation to fuller trophozoite forms and finally to schizonts that have replicated their DNA and formed daughter merozoites, the invasive forms that will invade new RBCs. We synchronized parasites within a three-hour window, washed out aTc in young rings, and monitored parasite development by thin smear. By 24 hours after invasion, Pf3D7_1437000-depleted parasites appeared smaller than their non-depleted counterparts (Fig. 6A, mean area +aTc = 6.95 μm^2^ vs. DMSO 5.45 μm^2^, unpaired t-test gives p < 0.0001). The disparity in size persisted throughout the intraerythrocytic development cycle, with Pf3D7_1437000-depleted parasites approximately half the size of non-depleted parasites by 36 hours (mean area = 22 μm^2^ vs. 12 μm^2^) (Fig. 6A). By 48 hours after invasion, non-depleted parasites had formed schizonts filled with clearly distinguishable daughter cells. Pf3D7_1437000-depleted parasites apparently remained trophozoites: some mature but without distinguishable daughter cells, others shrunken (Fig. 6B). Correspondingly Pf3D7_1437000-depleted parasites remained much smaller than non-depleted parasites (mean = 18 μm^2^vs. 30 μm^2^) (Fig. 6A). Thus, we conclude that Pf3D7_1437000 is important for parasite growth, and for completion of proper schizogony.

**Figure 6.**
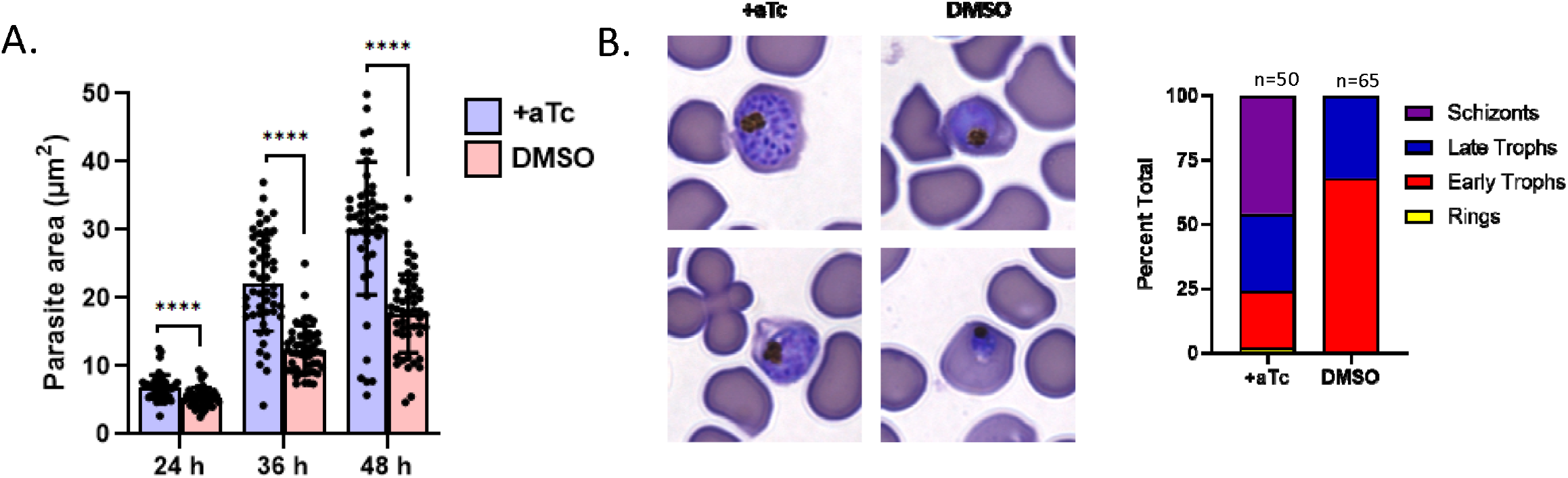
Pf3D7_1437000 depletion results in growth arrest before schizogony: (A) Area of parasites +/-aTc examined by thin smear at indicated times post-infection. Bar height indicates mean of 50 measured parasites, error bars indicate standard deviation. For each time point the measured areas were compared by an unpaired t-test, which yielded p-values < 0.0001 (marked with “****”). (B) At left, demonstrative images showing phenotypes described in the text. +aTc parasites were mostly schizonts and late trophozoites; parasites without aTc arrested before schizogony. At right, quantification of parasite life stages of 10 microscopic fields (n = 50 or 65 parasites, shown atop bar). Experiment was performed twice. A representative experiment is shown.

The Pf3D7_1437000 depletion phenotype appeared to us quite distinct from death due to protein export block, which arrests growth at the transition from rings to trophozoites (Beck et al., 2014). To distinguish these phenotypes more clearly, we fixed Pf3D7_1437000-regulated parasites alongside Hsp101-DD parasites at several time points during and assessed DNA content with the dye SYBR Green I by flow cytometry (Fig. 7). By 28 hours after invasion, Hsp101-disrupted parasites lag behind non-disrupted parasites, while Pf3D7_1437000-depleted parasites are indistinguishable from their non-depleted partners. By 40 hours after invasion Hsp101-disrupted parasites have the same DNA content that they did 20 hours earlier. Pf3D7_1437000-depleted parasites too lag behind their non-depleted counterparts, but with a substantially larger DNA content. Thus, it appears that growth arrest caused by Pf3D7_1437000 depletion is distinct from and substantially later than arrest caused by export disruption. Disrupting export prevents most of S-phase, while Pf3D7_1437000 depleted parasites continue until almost the final rounds of DNA replication.

**Figure 7.**
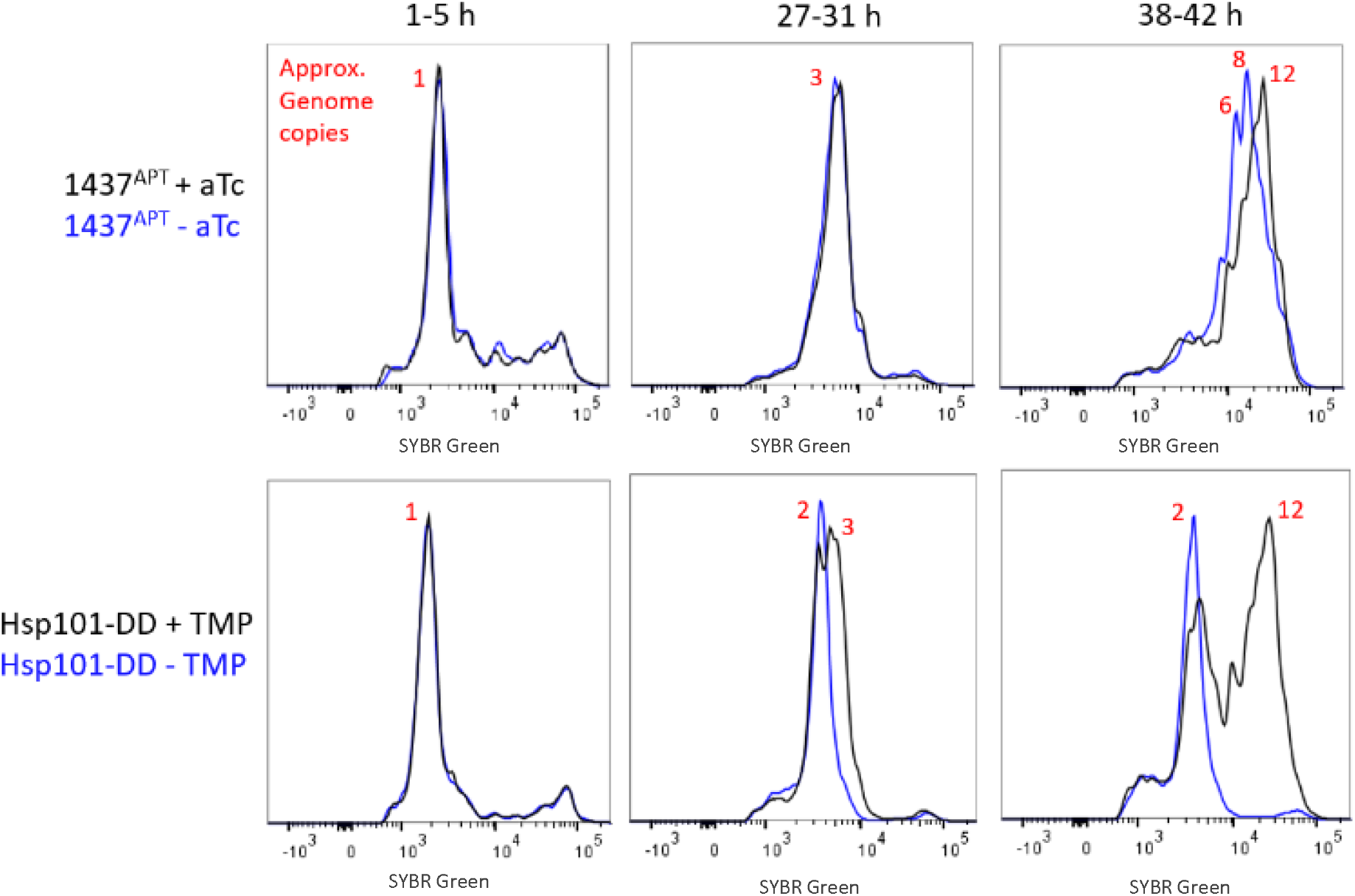
Pf3D7_1437000 depleted parasites do not complete DNA replication: Synchronized parasites were fixed at the indicated times, and their DNA content measured with SYBR Green I and flow cytometry. Samples were run in technical triplicate (N=100,000 cells per sample). Uninfected RBCs were gated out by their low SYBR Green I fluorescence. Histograms are shown, with SYBR Green I fluorescence along the x-axis, and frequency on the y-axis. Approximate genome copy numbers were estimated from each peak’s fluorescence intensity, assuming most parasites at 1 to 5 hours after invasion have a single copy of the genome. This experiment was performed twice; a representative experiment is shown.

## Discussion

Here we sought to investigate the acetylation of PEXEL proteins, instead finding a putative NAT that is essential for parasite growth but appears to be uninvolved in protein export and likely not the PEXEL NAT. Pf3D7_1437000 seemed a likely candidate (and really the only plausible candidate) to accomplish this post-PM V-cleavage acetylation. Indeed there is explicit speculation in the literature that it may serve such a role (Boddey and Cowman, 2013; Osborne et al., 2010). Supporting that is its putative ER localization: it appears in Marapana, et al.’s extensive immunoprecipitation/mass spectrometry results when pulling down SPC21 (signal peptidase) and SPC25 (Marapana et al., 2018). It has a clear *T. gondii* ortholog predicted to reside in the *Toxoplasma* ER (Barylyuk et al., 2020). Our findings are consistent with this ER localization, yet argue against a model where Pf3D7_1437000 acetylates PEXEL N-termini following PM V cleavage. The *Toxoplasma* PMV ortholog, ASP5, resides in and processes proteins in the Golgi, so N-acetylation after protein processing in this organism is not likely to involve the same NAT ortholog.

Our data most parsimoniously support a model where Pf3D7_1437000 is not the PEXEL NAT; however, our data cannot yet exclude alternative models. First, PF3D7_1437000 could be so efficient at acetylating its substrates that the small amount of it remaining after knockdown could be capable of acetylating all HRP2 and HRP3 N-termini that we could detect. We tried to minimize this possibility by harvesting schizonts for our mass spectrometry experiments, giving any acetylation phenotype as much time as possible to manifest. At the time of harvest, Pf3D7_1437000 knockdown parasites have been physically affected by the knockdown (as measured by parasite size) for at least 20 hours, suggesting that the remaining enzyme is not able to accomplish its full function for much of the life cycle. Still, the possibility remains that Pf3D7_1437000 is acetylating HRP2 and HRP3 in our assay, and that parasite growth problems are caused by a distinct second function of this protein that is more sensitive to knockdown. Second, it remains possible that Pf3D7_1437000 depletion affects HRP2/HRP3 acetylation, but we failed to detect the unacetylated forms due to their poor stability in the parasites, solubility in our purification system, or ionization for mass spectrometry. The only data we have to consider in this light is that Tarr, et al. has described mutants of the reporter fusion REX3^1-61^-GFP that are unacetylated (and not exported) and appear in similar abundance to the wild-type protein (Tarr et al., 2013). That is, acetylation of the PEXEL N-terminus is not universally required for protein stability and detection; however, we cannot exclude the possibility that such problems have frustrated our analysis here.

Regardless of whether Pf3D7_1437000 is the PEXEL NAT, we report here that its depletion does not affect protein export, and that – while Pf3D7_1437000 is essential for parasite development – the phenotype of its depletion differs from described disruptions of export machinery. Depletions/disruptions of the major PTEX components Hsp101 and PTEX150 have been described, each of which arrested parasite growth as early trophozoites (Beck et al., 2014; Elsworth et al., 2014). Chemical inhibition or depletion of PM V can arrest parasite growth immediately after invasion, or as early trophozoites (Boonyalai et al., 2018; Polino et al., 2020; Sleebs et al., 2014). Here we show that Pf3D7_1437000 depletion arrests parasite growth substantially later than we observe when blocking export via Hsp101 disruption. Our finding that Pf3D7_1437000 depletion affects parasite size through much of intraerythrocytic development cycle is curious, but its causes could be manifold, and further study of Pf3D7_1437000-depleted parasites would be needed to elucidate how this putative enzyme could be influencing cell size.

Among our field’s directives is to identify and characterize essential parasite proteins and pathways sufficiently diverged from their hosts’ orthologues as to be specifically targeted with chemotherapeutics. In some ways, Pf3D7_1437000 fits that bill: an essential putative NAT with orthologues throughout Apicomplexa, yet without a clear orthologue in mammals. However, hurdles remain before its druggability can be assessed. First, the degree to which this enzyme is essential in other pathogenic apicomplexans is not yet known. The *Plasmodium berghei* knockout screen scored the P. *berghei* orthologue PbANKA_0611800 as “essential” albeit with low confidence (Bushell et al., 2017), while the genome-wide CRISPR screen in *Toxoplasma* gondii assigned the likely orthologue TgME49_305450 to have an intermediate phenotype score, higher than most validated essential genes, and lower than most validated dispensable genes (Sidik et al., 2016). Delineating how well-conserved the function and essentiality of Pf3D7_1437000 is remains for future work. Perhaps more importantly – assuming Pf3D7_1437000’s essential function is as an NAT – chemical inhibition of NATs has only been described with acetyl-CoA-peptide conjugates (Foyn et al., 2013). The development of more druglike and cell permeant NAT inhibitors might raise the profile of this enzyme class for additional chemotherapeutic development.

Our data leave us with the question: if Pf3D7_1437000 is not the PEXEL NAT, what is? The simplest possibility is that another NAT resides in the parasite ER, with the most obvious candidates being the additional members of the GNAT superfamily listed in Figure 1. Each has an ortholog with an alternative described function in yeast and mammals, but perhaps they serve a distinct role in Apicomplexa. An alternative is that the genome continues to hide an as-yet cryptic NAT whose sequence defies our attempts to computationally predict its function. Unfortunately no obvious candidate jumps out from the Marapana, et al. immunoprecipitations of various ER proteins involved in PEXEL processing (Marapana et al., 2018). A third possibility is that no protein NAT truly resides in the parasite ER, but instead that following PEXEL cleavage by PM V, the new N-terminus somehow accesses the cytosol and is acetylated by the regular cadre of ribosome-associated NATs. Tarr, et al. used a split-GFP setup to show that the active site of PM V is oriented into the ER lumen (Tarr and Osborne, 2015), and we have since presumed that the post-cleavage PEXEL N-terminus is limited to the ER lumen; however, our understanding of the dynamics of secretory import and traffic in *Plasmodium* is itself limited, and perhaps peptides can somehow transiently access cytosolic NATs.

Lastly, the determinants of exported protein trafficking in *P. falciparum* remain unclear. Here, we were unable to assess the role of PEXEL acetylation on export competence. However, our mass spectrometry approach measured the mass of intact HRP2 and HRP3 and found each to be the exact mass expected from the polypeptide backbone and N-terminal acetylation alone, suggesting that export competent proteins need no further post-translational modification beyond what has already been described.

Taken together, our data provide new insight into the processing of exported proteins – if largely by excluding the primary candidate for involvement in the process. Additionally we provide initial characterization of the putative NAT Pf3D7_1437000, whose essential function in schizogony remains unclear. We hope that future work will uncover the identity and role of the PEXEL NAT, and also clarify the role of Pf3D7_1437000 in the *Plasmodium* life cycle.

## Acknowledgements

Our mass spectrometry approach was formed in consultation with and performed by the Donald Danforth Plant Science Center Proteomics & Mass Spectrometry Facility, particularly Brad Evans and Shin-Cheng Tzeng. Additionally we thank Geoffrey McFadden for the anti-ACP antibody, Dianne Taylor for anti-HRP2, John Adams for anti-ERD2 (through the Malaria Research and Reference Reagent Resource Center), Odile Mercereau-Puijalon for anti-FIKK4.2 (through the European Malaria Reagent Repository), and Joshua Beck for sharing the NF54^attB^ -DiCre parasite line, pM2GT-mNeonGreen-3xHA plasmid, and pM2GT-mRuby3-3xFLAG plasmids pre-publication.

## Funding

This work was supported by an American Heart Association Predoctoral Fellowship (#18PRE33960417, awarded to A.P.) and by the National Institute of Allergy and Infectious Diseases (RO1 AI047798, awarded to D.G.). The Donald Danforth Plant Science Center Proteomics and Mass Spectrometry Facility acknowledges the support of the National Science Foundation (#DBI-0922879) for acquisition of the LTQ-Velos Pro Orbitrap LC-MS/MS.

## Accession Numbers

For all *Plasmodium* proteins referenced in this study, the PlasmoDB accession numbers are below:

### Putative NATs

Pf3D7_0109500, Pf3D7_0629000, Pf3D7_0805400, Pf3D7_0823300, Pf3D7_1003300, Pf3D7_1020700, Pf3D7_1323300, and Pf3D7_1437000.

### Organellar markers

plasmepsin II (PF3D7_1408000), plasmepsin V (PF3D7_1323500), ERD2 (PF3D7_1353600), ACP (PF3D7_0208500), and aldolase (PF3D7_1444800).

### Export-related proteins proteins

Hsp101 (PF3D7_1116800) HRP2 (Pf3D7_0831800), FIKk4.2 (PF3D7_0424700), HRP3 (PF3D7_1372200).

### Discussion

SPC21 (PF3D7_1331300), SPC25 (PF3D7_0320700), REX3 (PF3D7_0936300), PTEX150 (PF3D7_1436300).

## Author contributions

AJP conceptualized this work, performed experiments, and wrote the manuscript under the supervision of DEG. YAC, KF, and YY performed experiments and influenced the path of this project by sharing their ideas.

## Materials and methods

### Generation of plasmids

The construct for regulating Pf3D7_1437000 levels was made using the previously described pSN054 vector (Nasamu et al., 2021; Polino et al., 2020). Primer sequences are listed in Supplementary Table 1; shorthand names will be used here. Homologous sequences for genome repair were amplified from NF54^attB^ (Nkrumah et al., 2006) with primers 14APT-1/14APT-2 for the left/upstream homologous region, and 14APT-3/14APT-4 for the right/downstream homologous region. The resulting PCR products were inserted into pSN054 cut with FseI and I-SceI respectively via Gibson Assembly (New England Biolabs). The recodonized Pf3D7_1437000 gene was synthesized by GENEWIZ (South Plainfield, NJ) and inserted into pSN054 containing the above homologous regions. Plasmid was cut with AsiSI and the recodonized gene inserted via Gibson Assembly, to make the final vector called pSN054-1437000-3xHA. The vector was transformed into BigEasy-TSA Electrocompetent Cells (Lucigen) for propagation. When amplifying vector for harvest, 0.01% weight/volume arabinose was added to stimulate plasmid replication.

The two endogenous tagging vectors used for live microscopy: pM2GT-1437000-mNeonGreen-3xHA (yDHOD) and pM2GT-PMV-mRuby3-3xFLAG (hDHFR) are derived from pM2GT-EXP2-mNeonGreen (yDHOD) (Glushakova et al., 2017) and pM2GT-Hsp101-3xFLAG (Garten et al., 2018; Ho et al., 2018). For the former, mNeonGreen-3xHA was amplified (and the HA added) by primers NG-HA-F/NG-HA-R. The amplicon was inserted into pM2GT-EXP2-mNeonGreen cut with AvrII/EagI via In-Fusion Cloning. For tagging Pf3D7_1437000, homologous sequences were amplified from the NF54^attB^ genome with primers 14NG-1/14NG-2 for the right homologous region (in the 3’ UTR), and 14NG-3/14NG-4 for the left homologous region. The two PCR products were combined in an overlap-extension PCR with the right homologous region forward and left homologous region reverse primers, and the resulting PCR product gel extracted and inserted into the XhoI/NheI-cut plasmid via In-Fusion Cloning (Takara) to make the donor vector pM2GT-1437000-mNeonGreen-3xHA, which was then transformed into XL10-Gold Ultracompetent Cells (Stratagene) for propagation.

Synthesis of the PM V tagging vector was analogous. First mRuby3 was amplified with primers Rub-H-F/Rub-H-R and added to pM2GT-Hsp101-3xFLAG (hDHFR) (Garten et al., 2018; Ho et al., 2018) cut with AvrII using In-Fusion Cloning. To adapt the plasmid to PM V tagging we used primers PMVR-1/PMVR-2 and PMVR-3/PMVR-4 to amplify the right and left homologous regions respectively. These PCR products were inserted into the Xho/NheI-cut plasmid in a single pot reaction with In-Fusion Cloning, and the resulting vector transformed into XL10-Gold cells for propagation.

CRIPSR/Cas9 targeting plasmids for each were made in the previously described pAIO3 plasmid (Nessel et al., 2020). Primers 14G-1, 14G-2, 14G-3, 14G-4, PMVG-1, and PMVG-2 were each ordered along with their reverse complement sequences, annealed in the thermal cycler, and then inserted into AvrII-cut pAIO3 by In-Fusion Cloning. Completed vectors were transformed into XL10 Gold cells for propagation.

### Parasite culture

For all experiments described here we cultured *P. falciparum* in PRMI 1640 (Gibco) supplemented with 0.25% (w/v) Albumax I, 15 mg/L hypoxanthine, 110 mg/L sodium pyruvate, 1.19 g/L HEPES, 2.52 g/L sodium bicarbonate, 2 g/L glucose and 10 mg/L gentamycin with human RBCs added to 2% hematocrit. Parasite cultures were maintained in sealed chambers under a gas mixture consisting of 5% O_2_, 5% CO_2_, and 90% N_2_ at 37°C. Deidentified RBCs were obtained from Barnes-Jewish Hospital blood bank (St. Louis, MO), St. Louis Children’s Hospital blood bank (St. Louis, MO), and the American National Red Cross.

### Generation of parasite lines

All genetic modifications described here were done in the parasite line NF54^AttB^ (Nkrumah et al., 2006) referred to as “NF54” throughout. For each transfection, donor vectors were linearized if necessary (pSN054 is already linear) and 50 μg each of donor vector and pAIO3 with relevant guide were combined, ethanol precipitated to ensure sterility, and dissolved in 100 μL sterile water. At time of transfection, the dissolved DNA was brought up to 400 μL in cytomix (120 mM KCl, 0.15 mM CaCl_2_, 2 mM EGTA, 5 mM MgCl_2_, 10 mM K_2_HPO_4_, and 25 mM HEPES adjusted to pH 7.6 with KOH; plasmid is solubilized more effectively in water than cytomix, so we typically allow DNA to dissolve in 100 μL water, then when dissolved add 100 μL 2x cytomix and 200 μL 1x cytomix to bring it up to transfection volume) and transfected into ∼5% young ring-stage parasites with a BioRad Gene Pulser II (Settings: 0.31 kV, 0.950 μF, capacitance set to “High Cap”, resistance on the Pulse Controller II set to “Infinite”). Successful transfectants were selected with the relevant drug beginning 24 hours after transfection: Blasticidin S (2.5 μg/mL; Fisher) for the aptamer line, DSM-1 (2 μM; Asinex) and WR-99210 (5 nM; gift from D. Jacobus of Jacobus Pharmaceutical Co.) for the mNeonGreen- and mRuby3-tagged line. Medium was changed daily for the week following transfection, then thrice weekly until parasites could be visualized by thin smear, typically two to four weeks after transfection.

### Validation of lines

Proper integration of the pSN054-1437000-3xHA vector was confirmed by Southern blotting. Genomic DNA from the parent and edited lines was isolated (QIAamp DNA Blood Miniprep Kit) and 10 μg each digested with HinDIII, separated overnight on a 0.7% agarose gel, and transferred to nylon (Nytran SuPerCharge TurboBlotter, 0.45 μm, GE) overnight. The blot was then probed with the left homologous region (PCR product of 14APT-1/14APT-2) labelled with alkaline phosphatase (Amersham AlkPhos Direct Labelling Kit; GE) in hybridization buffer (Amerhsam) at 55°C overnight, washed twice each in primary wash buffer (120 g/L urea, 1 g/L SDS, 100 mL/L 0.5 M sodium phosphate pH 7, 8.7 g/L NaCl, 2 g/L Amersham blocking reagent) and secondary wash buffer (6.05 g/L Tris base, 5.6 g/L NaCl, 2 mL/L 1M MgCl_2_, pH 10), then detected with Amersham CDP-Star Detection Reagent (GE) and exposed to blue autoradiography film (MidSci, BX810) overnight.

Additional validation was by western blot done exactly as described in (Polino et al., 2020) (section “Validation of PMV^APT^ line”).

### Assessment of Pf3D7_1437000 depletion

In all experiments shown, infection with the aTc-regulatable line (1437^APT^) was synchronized by purifying schizonts on LD columns (Miltenyi Biotech), eluting into fresh blood and media, lacking aTc and allowing parasites to invade for 3-4 hours. Invasion was halted by replacing media with 5% sorbitol, lysing any un-egressed schizonts. We then washed in fresh media one additional time for five minutes, to ensure aTc was removed from the culture. These synchronized parasites were cultured in either the presence of 100 nM aTc (“+aTc” throughout) or an equal volume of DMSO (“DMSO” throughout). DiCre excision experiments were performed as above, but parasites were cultured in the presence of 100 nM aTc and either 50 nM rapamycin or an equal volume of DMSO.

### Western blotting

We performed western blotting as in (Polino et al., 2020) with primary antibodies mouse anti-HA diluted 1:1000 (clone 16B12; Biolegend #901501), rabbit anti-PfAldolase diluted 1:2000 (abcam #ab207494, targets the protein with PlasmoDB accession PF3D7_1444800), and mouse anti-FLAG diluted 1:500 (Sigma #F1804), followed by secondary antibodies goat anti-mouse IRDye 800CW (Licor) and donkey anti-rabbit IRDye 680RD (Licor) both diluted 1:10,000. For Figure 2, panel C parasites were harvested at the indicated times and Pf3D7_1437000-3xHA levels quantified using ImageStudio Lite v. 5.2 (Licor). The sizes of bands were approximated using the Precision Plus Protein Dual Color Standards (Bio-Rad; #1610374).

### Flow cytometery

To assess the effect of Pf3D7_1437000 depletion on parasite growth parasites were maintained in technical triplicate (3 × 1mL culture) and their growth monitored daily by flow cytometry (BD FACSCanto with attached High Throughput Sampler) by diluting culture 1:20 into PBS with 0.8 μg/mL acridine orange (Molecular Probes).

We assessed parasite progress through the DNA replication cycle as in (Perrin et al., 2021). Briefly, parasites were synchronized as above, then fixed at the indicated times (see Figure 7) by doubling their volume in 2x PBS + 0.4% glutaraldehyde (final concentration, 1x PBS + 0.2% glutaraldehyde) and stored at 4°C until all time points collected. Then DNA was stained using SYBR Green I (ThermoFisher), measured on a BD FACSCanto, and analyzed with FloJo v. 10.7.1.

### Microscopy

For localization of Pf3D7_1437000, parasites were synchronized as above to within a 4-hour window, then harvested 24 hours after invasion ended (i.e. parasites 24-28 hours old), washed once in PBS, and prepared for immunofluorescence imaging as recommended in (Tonkin et al., 2004) with the modification that cells were settled onto concanavalin A-coated coverslips (0.5 mg/mL) for 10 minutes prior to fixation, and that the wash following primary and secondary antibody incubation consisted of five PBS washes for three minutes each. Primary antibodies: mouse anti-HA (close 16B12, Biolegend #901501) diluted 1:100, rabbit anti-HA (Sigma-Aldrich, H6908) diluted 1:100, mouse anti-PMV (Banerjee et al., 2002) diluted 1:50, rabbit anti-ACP (Waller et al., 1998) diluted 1:100, and rabbit anti-aldolase (abcam #ab207494) diluted 1:500. Secondary antibodies: goat anti-mouse IgG-AlexaFluor488, goat anti-rabbit IgG-AlexaFluor555 (both from Invitrogen) diluted 1:2000. Coverslips were mounted in ProLong Gold Antifade with DAPI (ThermoFisher), allowed to cure for 24 hours, then imaged on a Zeiss AxioImager.M1 epifluorescence microscope with a Hamamatsu ORCA-ER CCD camera and AxioVision v. 4.8.1. Images were cropped, scale bars added, and brightness/contrast adjusted for presentation with Zen Lite v. 2.5 (Zeiss).

Parasites for live microscopy were synchronized as in the preceding paragraph, harvested 24 hours after invasion, washed once in PBS, incubated with 1 μg/μL Hoechst 33342 (Invitrogen; H3570) for 30 minutes, then imaged on the same AxioImager.M1 described above. Images were analyzed in ImageJ: regions-of-interest were drawn around the mNeonGreen signal by hand, applied to all channels, and Pearson’s Correlation Coefficients calculated using ImageJ’s coloc.pearsons() function.

To monitor parasite size, 1437^APT^ was synchronized as above and monitored by thin smear at the indicated times (see Figure 6). Thin smears were fixed and stained with Harleco Hemacolor (MilliporeSigma), then imaged using a Zeiss Axio Observer.D1 at the Washington University Molecular Microbiology Imaging Facility. Parasite size was assessed using ImageJ, by manually drawing parasite borders and calculating their area.

To assess the effect of Pf3D7_1437000 depletion on protein export, 1437^APT^ was synchronized as above alongside Hsp101-DD (Beck et al., 2014), with Hsp101-DD maintained in 10 μM trimethoprim (TMP) or an equal volume of DMSO. Parasites were harvested 28 hours after the invasion ended (i.e. parasites were 28-32 hours old), fixed, and processed as above. Primary antibodies are mouse anti-HRP2 clone 2G12 (Rock et al., 1987) diluted 1:500, mouse anti-FIKK4.2 (Kats et al., 2014) diluted 1:500, and mouse anti-KARHP clone 18.2 diluted 1:500.

### Sorbitol lysis assay

To measure sensitivity to sorbitol lysis, parasites were maintained as in the preceding paragraph. 28 hours after invasion, parasites were moved to a 96-well plate (100 μL/well), pelleted, and the media supernatant aspirated off. Every 5 minutes, we resuspended another row of the plate (4 conditions, 3 technical replicates per condition) in 5% sorbitol. One row was instead resuspended in deionized water to fully lyse the RBCs. After 24 minutes, the infected RBCs were again pelleted and the supernatant transferred to a new 96-well plate. Lysis of infected RBCs was measured by absorbance at 405 nm, a measure of hemoglobin abundance, using an Envision Multimodal Plate Reader (PerkinElmer). Values are expressed relative to the deionized water control (representing 100% lysis). The cultures were at 6% parasitemia at the outset of the experiment, so a maximum expected value is 6% lysis.

### Assessment of HRP2/3 acetylation

To investigate the acetylation status of HRP2 and HRP3, 1437^APT^ parasites were synchronized and maintained as above. 44 hours after invasion, aTc was washed from cultures (3 washes; 5 min. each), and cultures resuspended in 500 nM aTc or an equal volume of DMSO. For the DiCre experiments, either 50 nM rapamycin or an equivalent volume of DMSO were added. At the end of the following cycle (again, 44 hours after inivasion) parasites were pelleted, media was aspirated off, and then the RBC and PV lysed in 0.035% saponin in PBS at 4°C for 15 minutes. Parasite material was then pelleted at 17,000 g and the supernatant poured onto nickel HTC agarose beads (Gold Biotechnology) that had been equilibrated in PBS + 100 mM imidazole. Columns were then washed in 20 column volumes of PBS + 100 mM imidazole to remove the abundant hemoglobin, then eluted in 15 mL PBS + 1 M imidazole. Eluate was concentrated in 10K cutoff Amicon Ultra-15 Centrifugal Filters (Millipore), flash frozen in liquid nitrogen, then sent to the Danforth Plant Science Cetnter’s Proteomics and Mass Spectrometry Facility.

There, samples were acidified by formic acid to 1% then cleaned up with C4 ZipTip (Millipore). The captured samples were eluted with 50% acetonitrile in 0.1% formic acid, dried down and then resuspended in 10 μL 3% acetonitrile in 1% formic acid. 5 μL of sample was analyzed by LCMS with a Dionex RSLCnano HPLC coupled to an OrbiTrap Fusion Lumos (Thermo Fisher Scientific) mass spectrometer using a 60 min gradient (2-90% acetonitrile). Sample was resolved using a 75 μm x 150 cm PepMap C4 column (Thermo Scientific). MS spectra of protein ions of different charge-states were acquired in positive ion mode with a resolution setting of 120,000 (at m/z 200) and accurate mass was deconvoluted using Xcalibur (Thermo Scientific).

## Supplementary Figures

**Supp. Fig. 1.**
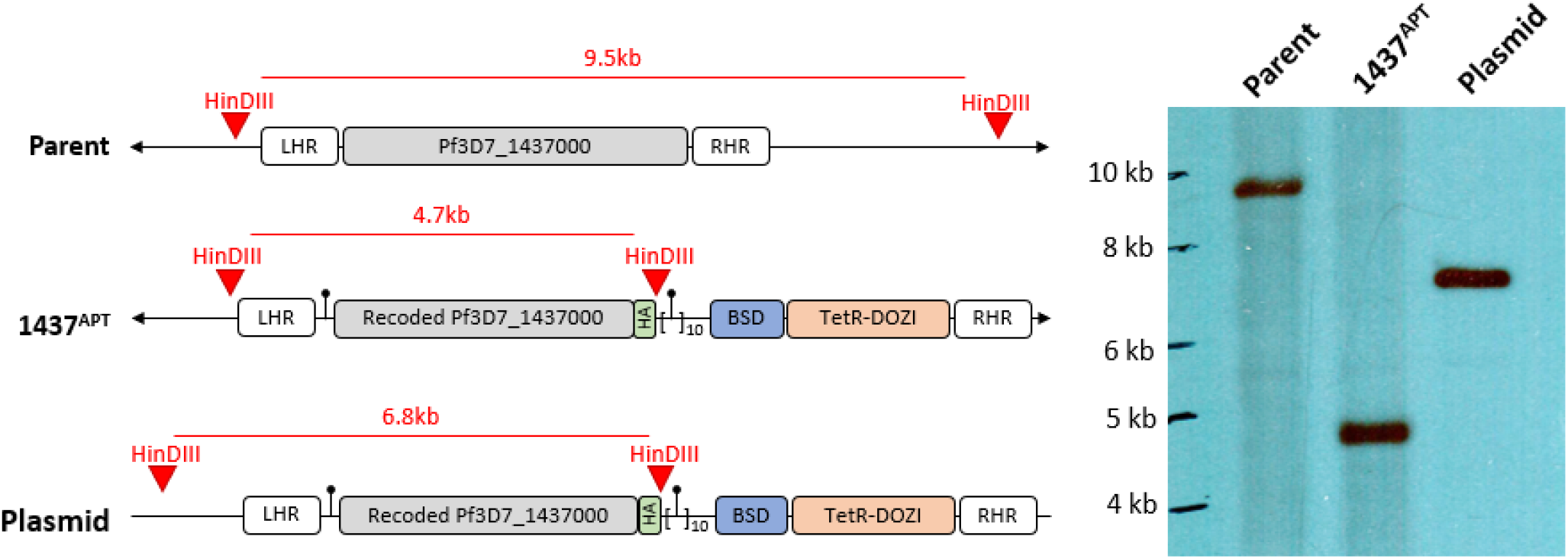
Southern blot: Correct genome editing was verified by Southern blot with left homologous region (LHR) used to probe the HinDIII-fragmented genome. Digest schematic shows expected size of bands for the parent, edited line (1437^APT^) and donor plasmid.

**Supp. Fig. 2.**
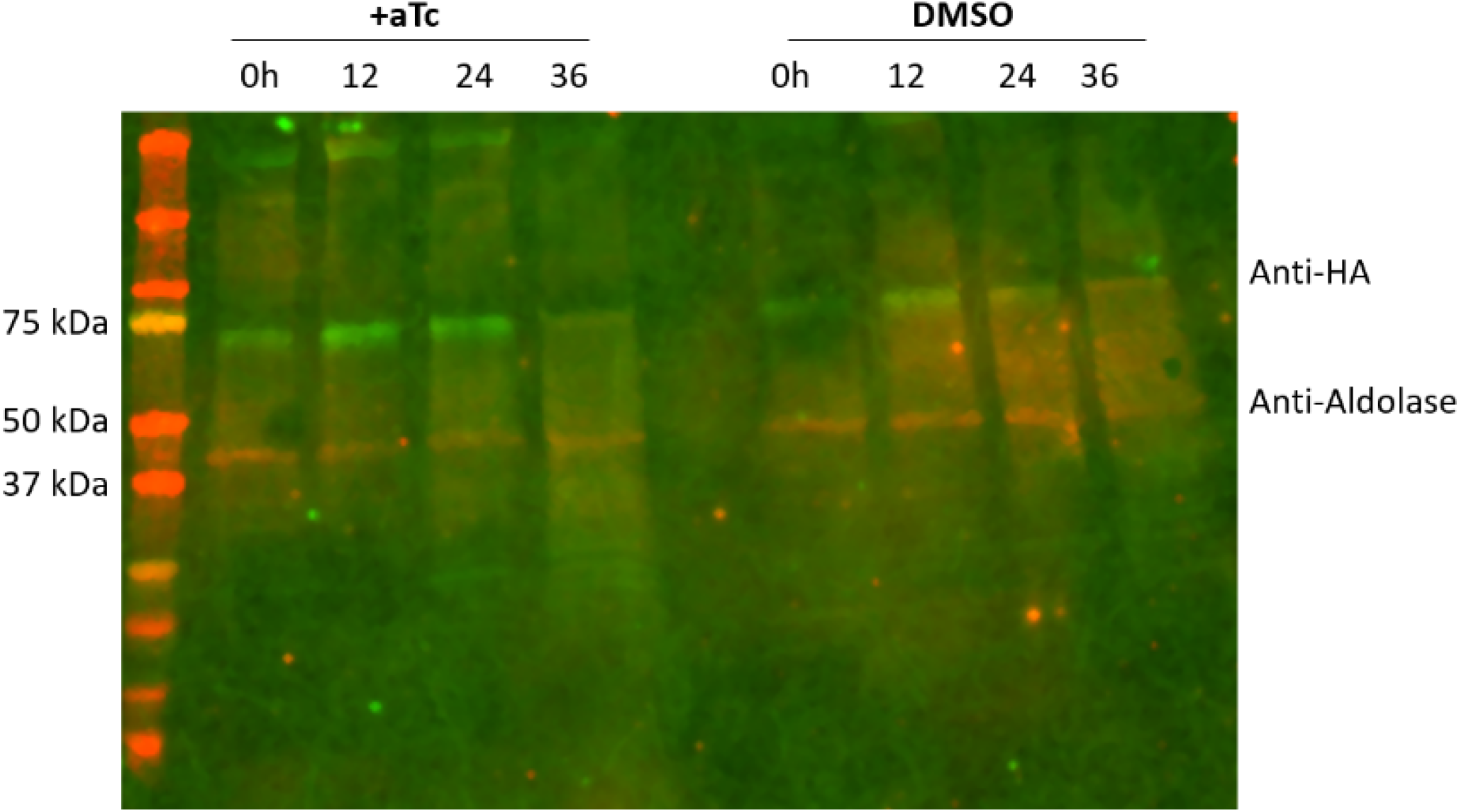
Uncut gel for Figure 1.

**Supp. Fig. 3.**
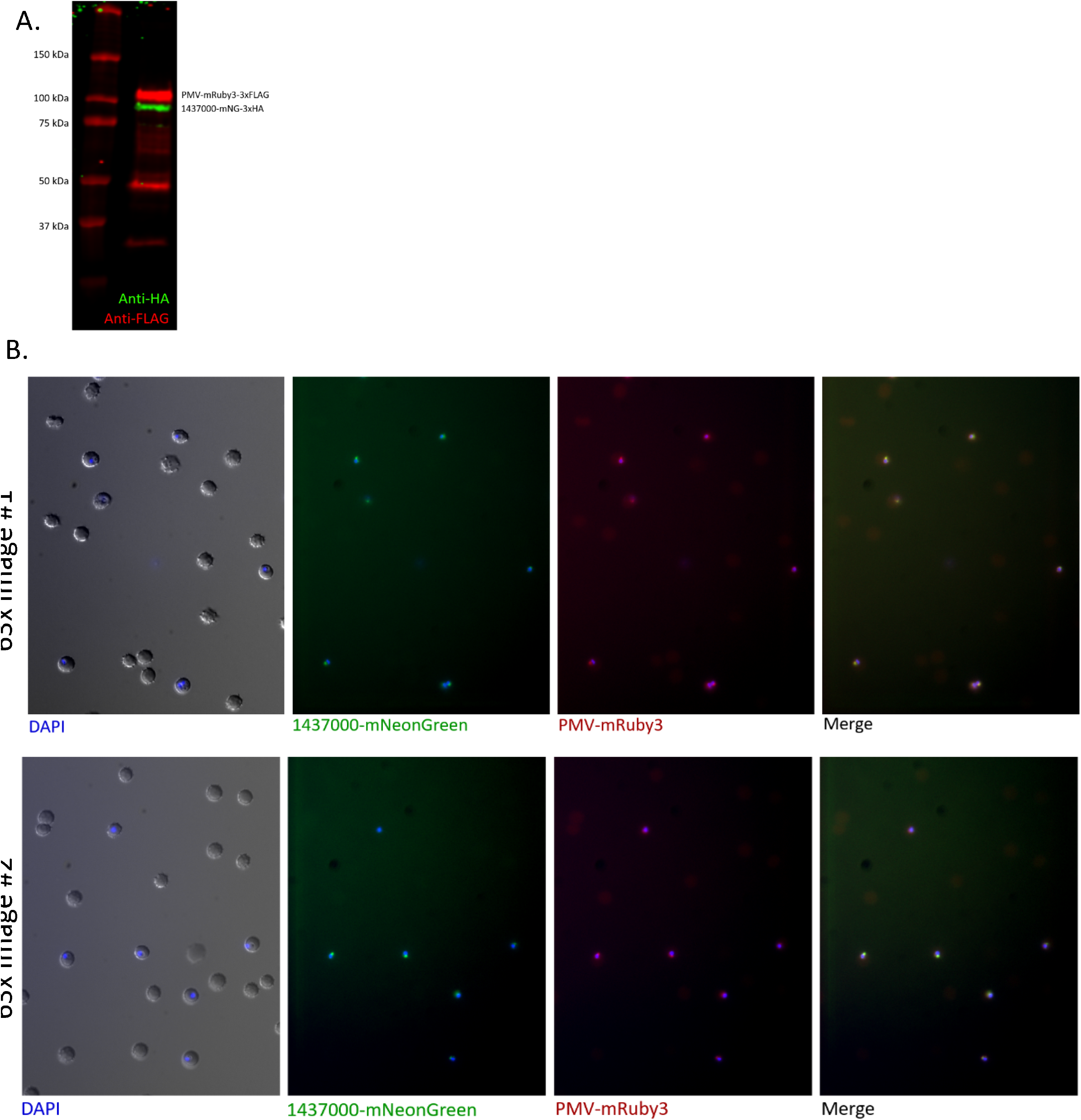
Supplement to Figure 3B: (A) Western blot probing Pf3D7_1437000-mNeonGreen-3xHA and PM V-mRuby3-3xFLAG double-tagged line with antibodies for anti-HA (green) and anti-FLAG (red). (B) Additional images from Fig. 3B.

**Supp. Fig. 4.**
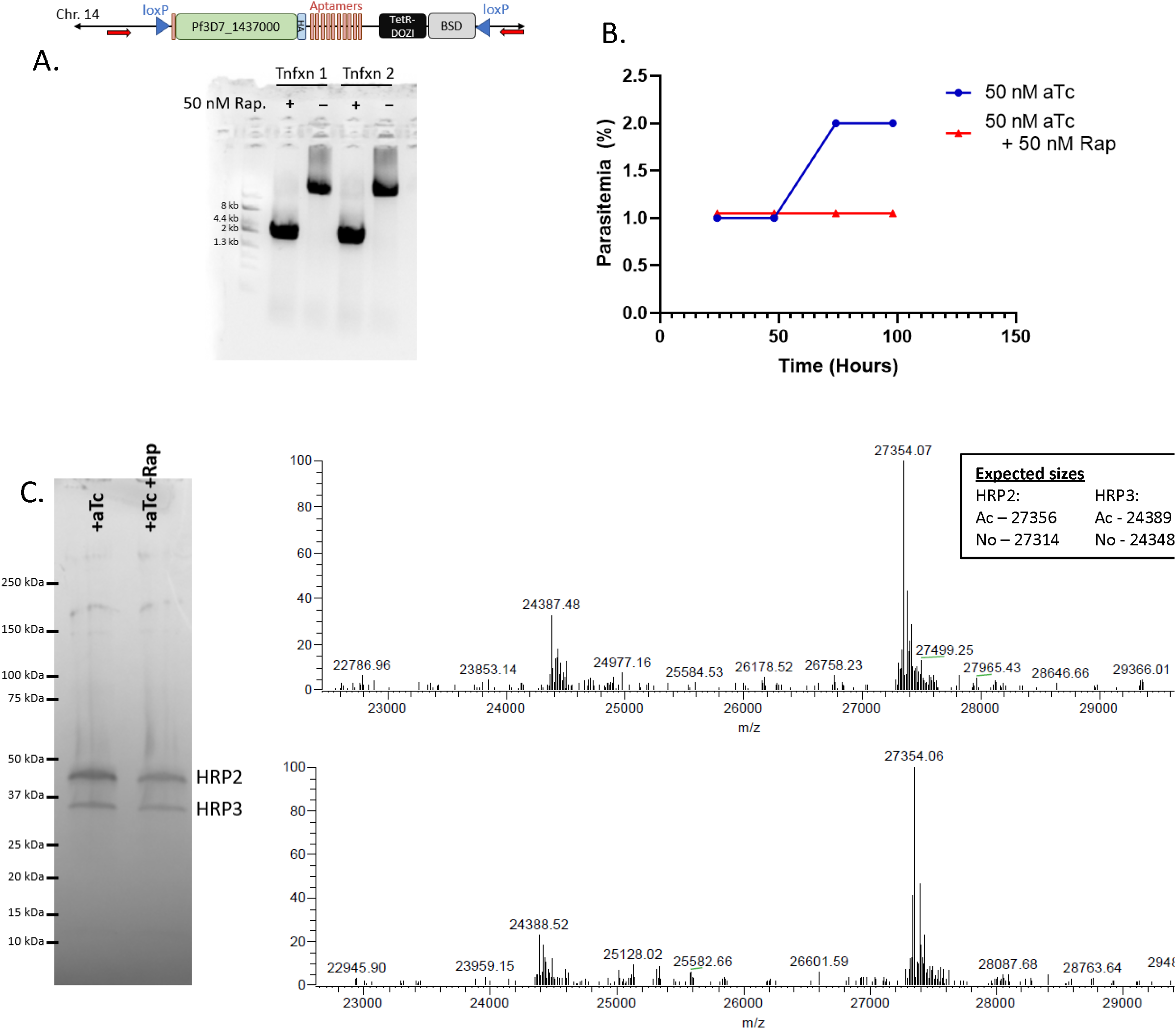
DiCre excision of Pf3D7_1437000 locus also doesn’t affect HRP2/HRP3 acetylation. (A) PCR to check excision of Pf3D7_1437000 locus upon addition of rapamycin (Rap) to cultures. At top, red arrows represent PCR primer locations. Expected sizes are 11 kb for the parent locus, 4 kb for the excised locus. Below, an ethidium bromide-stained agarose gel showing PCR products 24 hours after excision from two different transfections (“Tnfxn1” and “Tnfxn2”). (B) Rapamycin was added to ring stage parasites and growth followed daily by flow cytometry. Experiment was performed twice; a representative experiment shown. (C) Deconvoluted mass spectra from analysis of intact HRP2 and HRP3. Inset shows anticipated sizes for acetylated and un-acetylated HRP2 and HRP3 after PEXEL cleavage. This experiment was performed twice; a representative experiment is shown.

**Supp. Table 1:**
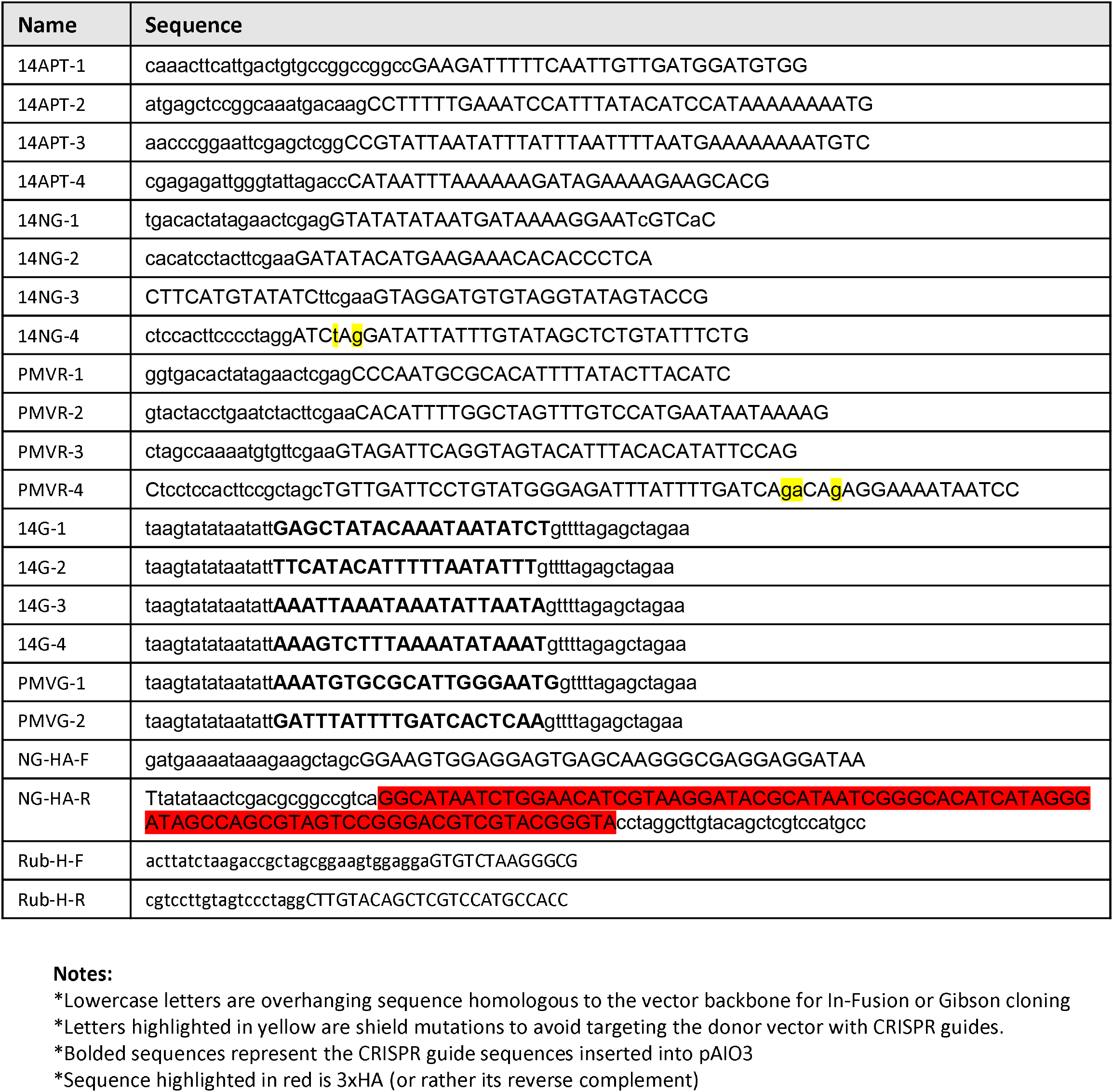
Primers used in this study.

## Notes

### Competing Interest Statement

The authors have declared no competing interest.

